# Instrumental Punishment Learning Enhances the Intrinsic Excitability of Basolateral Amygdala Neurons

**DOI:** 10.1101/2025.05.02.651812

**Authors:** Eddie T Wise, Philip Jean-Richard-dit-Bressel, Joanna O-Y Yau, Gavan P McNally, John M Power

## Abstract

The basolateral amygdala (BLA) is a structure that is critical for forming Pavlovian and instrumental emotional associations. Previous studies have established that Pavlovian fear conditioning increases the intrinsic excitability of BLA projection neurons. Learning-associated increases in excitability are hypothesised to facilitate the induction of synaptic plasticity and integration of neurons into functional learning circuits. Here, we examined whether instrumental learning produces similar changes in intrinsic excitability. Male and Female Long Evans rats were trained to lever press for a food reward and subsequently split into 3 behavioural groups; punished, yoked, and reward. Punished rats received response-contingent footshocks, yoked rats received response-independent footshocks, and reward rats received no shock. Intrinsic excitability was measured as the number of action potentials (APs) evoked by a series of depolarising current injections during whole-cell patch clamp recordings. Reward learning increased the intrinsic excitability of BLA projection neurons compared to naive controls. Punishment training maintained excitability and excitability was correlated with performance on the punishment task. Response-independent footshocks arrested reward-associated increases in excitability, reverting the BLA neurons to a naive state. Group differences were observed in AP half-width and the current underlying the post-burst AHP. Together, these findings show that instrumental reward learning enhances the intrinsic excitability of BLA projection neurons, potentially supporting action-outcome encoding by promoting synaptic plasticity and circuit integration. The results also suggest that different forms of learning engage distinct biophysical adaptations. A mechanistic understanding of the neurobiology underlying instrumental association learning may help inform the development of treatments for psychiatric disorders characterized by deficits in this process, such as major depressive disorder and conduct disorder.

## Introduction

An essential component of adaptive behaviour is the ability to evaluate the consequences of one’s actions. Through experience, animals and humans learn to associate specific behaviours with rewarding or punishing outcomes, allowing them to modify future behaviour to obtain reward and avoid punishment. This instrumental learning is critical for flexible, goal-directed behaviour and relies on the brain’s ability to form and store emotional associations (Ostlund & Balleine, 2008) and may be disrupted in disorders such as conduct disorder and addiction that can involve a failure to adapt behaviour despite adverse consequences (McNally et al., 2023). The basolateral amygdala (BLA) is a critical brain region for forming instrumental and Pavlovian emotional associations (Gottfried et al., 2002; Jean-Richard-Dit-Bressel et al., 2018; LaBar et al., 1998; Wassum & Izquierdo, 2015). Despite clinical relevance of instrumental emotional association learning, our knowledge of how the amygdala encodes emotional associations is largely restricted to Pavlovian conditioning.

Pavlovian conditioning is associated with several long-term neurophysiological changes to BLA neuronal circuits including long-term enhancement of excitatory inputs (Lynch, 2004), decreased local circuit inhibition (Lucas et al., 2016; Redondo et al., 2014), and enhanced intrinsic neuronal excitability (Motanis et al., 2014; Sehgal et al., 2023; Sehgal et al., 2014). While synaptic plasticity has long been the focus of most studies, there is growing recognition that intrinsic neuronal excitability, a neuron’s likelihood of firing in response to input, also plays a vital role in memory formation (Schulz, 2006; Sehgal et al., 2018). Changes in intrinsic excitability influence how neurons integrate synaptic signals and modulates the induction of synaptic plasticity (Power et al., 2011; Sah & Bekkers, 1996). Furthermore, neurons with elevated excitability are more likely to be incorporated into memory traces (Josselyn & Frankland, 2018; Josselyn & Tonegawa, 2020).

Although Pavlovian learning has been shown to increase intrinsic excitability in BLA neurons, it remains unclear whether similar changes occur following instrumental learning, particularly in response to punishment. Given the behavioural and clinical importance of instrumental learning, identifying whether distinct learning modalities induce distinct patterns of excitability in the BLA could reveal new principles of emotional memory encoding. Here, we examined whether instrumental training with either appetitive (reward) or aversive (punishment) outcomes alters the intrinsic excitability of principal neurons in the BLA. This work seeks to bridge the gap between Pavlovian and instrumental learning at the cellular level and to expand our understanding of how emotionally salient experiences shape intrinsic properties of neurons involved in emotional memory.

## Methods

### Subjects

Experiments were performed using 2-4-month-old male and female Long Evans rats (School of Psychology, UNSW). Rats were housed in groups of 2 to 4 in a colony room maintained on a 12:12 light dark cycle (lights on at 7:00). All procedures were approved by approved by the UNSW Animal Care and Ethics Committee and performed in accordance with the Animal Research Act 1985 (NSW), under the guidelines of the National Health and Medical Research Council Code for the Care and Use of Animals for Scientific Purposes in Australia (2013).

### Behavioural Training Procedures

Punishment training was performed as previously described (Jean-Richard-Dit-Bressel et al., 2022). All behavioural procedures were conducted in a set of eight identical experimental chambers [24 cm (length) × 30 cm (width) × 21 cm (height); Med Associates Inc.]. The floors consisted of steel rods (4 mm diameter, 15 mm apart) connected to a constant-current generator to deliver footshocks. The centre of the right side-wall included a recess (5 × 3 × 15cm) that housed a magazine dish (3 cm diameter) into which 45 mg grain pellets (Bio-Serv, NJ, USA) were delivered. The magazine was flanked by two retractable levers (2 × 4 cm) on either side of the magazine dish. All chambers were connected to a computer with Med-PC IV software (Med Associates Inc.), which controlled all required manipulandum and recorded all required variables.

Rats were food restricted for one day in their home cages and then received one experimental session of lever press acquisition for food reward per day. Naive animals were given additional food to match what was earned during the behavioural task. Body weights were maintained within 90% of the weight at the commencement of training and individual food intake was not monitored. Across training sessions, rats received daily access to 5-10 g food post-training and unrestricted daily access to water in their home cages. Initially, they were placed in the experimental chambers for two one-hour magazine training sessions, where lever-pressing on either of two simultaneously presented levers (left and right) was reinforced with grain pellets on a fixed ratio-1 (FR-1) schedule. In these initial sessions a lever retracted once it was pressed 25 times or at the cessation of the one-hour session. In subsequent training sessions, levers were then presented individually in an alternating pattern so that one lever was extended for 5 min while the other lever was retracted. After 5 min, the extended lever was retracted and the retracted lever was extended, such that each lever was always presented on its own. This alternation occurred throughout the 40-min session for a total of eight trials, four for each lever. Lever-pressing on either lever was reinforced with pellets on a 30-sec variable interval (VI30) schedule. Rats received a total of seven sessions of VI30 lever press acquisition.

On VI30 Day 8, rats were randomly assigned into one of three behavioural conditions. Rats in the “Punished” group were given an identical session of lever press training, except responses on one lever (punished lever, R1) also delivered a 0.5 s, 0.4 mA footshock on an FR10 schedule (independent of pellet outcome). If a response was due to deliver both pellet and shock then both were delivered. Responses on the other lever remained unpunished (unpunished lever, R2). Each punished animal was paired with a corresponding “Yoked” animal that received the same number of shocks at the same time as the corresponding punished rat, but these shocks occurred independently of any instrumental action taken by the animal. So, the Punished rats were subject to an instrumental punishment contingency whereas Yoked rats undergo non-contingent Pavlovian fear conditioning. Rats allocated to a “Reward Learning” group received an additional session of VI30 lever press training and were not shocked during training.

### Behavioural Analysis

Behavioural measures, including lever presses, magazine entries, and shocks delivered were recorded using MedPC (Med Associates Inc.). Lever preference on VI30 Day 8 was computed as a preference ratio (P-ratio) per subject in the form A/(A+B), where A and B were total presses on R1 and R2 levers, respectively. Ratios of 0.5 indicate no difference in pressing between the two levers, ratios less than 0.5 indicate suppression of A relative to B, and ratios greater than 0.5 indicate elevation of A relative to B. These data were analysed via one-way between subject ANOVA.

To assess changes in responding per lever from VI30 day 7 to day 8, a training ratio (T-ratio) was computed in the form T = D8/(D7+D8), where D7 and D8 were the responses on each lever at day 7 and day 8 respectively. This provided a self-normalized quantification of changes to punished and unpunished lever-pressing on day 8 relative to day 7, where T ratios greater than 0.5 indicate an increase in pressing from day 7 to day 8, while T ratios less than 0.5 indicate a decrease in pressing. T_Pun_ reflects the change in preference for the punished lever, while T_Unpun_ reflects the change in preference for the unpunished lever. Due to spurious lever preferences before day 8, T_Difference_ (T_Unpun_ -T_Pun_) was used as the primary index of punishment learning in this study. Rats with T_Difference_ > 0 were deemed to be learners because they demonstrated greater suppression of punished relative to unpunished lever-pressing.

### Slice Preparation

Brain slices were prepared within 1 hour of the final behavioural session. Rats were deeply anesthetised with 5% isofluorane gas and decapitated. Brains were rapidly extracted and placed in ice-cold cutting neuroprotective artificial cerebrospinal fluid (ACSF) composed of N-Methyl-D-glucamine (NMDG) (95 mM), KCl (2.5 mM), NaH_2_PO_4_ (1.2 mM), NaHCO_3_ (30 mM), HEPES (20 mM), glucose (30 mM), ascorbic acid (5 mM), Thiourea (2 mM), N-Acetylcysteine (10 mM), CaCl_2_ (0.5 mM), and MgSO_4_ (10 mM) for 3-4 minutes. Coronal brain slices (300 μm) that included the BLA were prepared using a vibratome (Model VT1200, Leica, Wetzlar, Germany) and incubated for 10 mins in a 30°C recovery ACSF solution. Slices were then transferred to Braincubator (Payo Scientific, Sydney, Australia) where they were maintained at room temperature in holding ACSF solution containing NaCl (95 mM), KCl (2.5 mM), NaH_2_PO_4_ (1.2 mM), NaHCO_3_ (30 mM), HEPES (15 mM), glucose (20 mM), Ascorbic Acid (5 mM), Thiourea (2 mM), CaCl_2_ (2 mM), MgSO_4_ (2 mM) for one hour prior to the commencement of recordings. All solutions were pH adjusted to 7.3-7.4 with HCl or NaOH and gassed with carbogen (95% O_2_, 5% CO_2_).

### Electrophysiological Recordings

Brain slices were transferred to a recording chamber, maintained at 30°C and continuously perfused with oxygenated recording ACSF containing NaCl (124 mM), KCl (3 mM), NaH_2_PO_4_ (1.2 mM), NaHCO_3_ (26 mM), glucose (10 mM), MgCl_2_ (1.3 mM), CaCl_2_ (2.5 mM). Targeted somatic whole cell patch clamp recordings were made using a microscope (Zeiss Axio Examiner D1) equipped with 20x water immersion objective (1.0 NA), LED fluorescence illumination system (pE-2, CoolLED) and an EMCCD camera (iXon+, Andor Technology). Bath application of CNQX (20 μM) was applied during cell recordings to block background glutamatergic activity. Patch pipettes (3-5 MΩ) were fabricated from borosilicate glass using a two-stage vertical puller (PC-10; Narishige) and filled with an internal solution containing potassium gluconate (130 mM), KCl (10 mM), HEPES (10 mM), Mg_2_-ATP (4 mM), Na_3_-GTP (0.3 mM), EGTA (0.3 mM), and phosphocreatine disodium salt (10 mM) (pH adjusted to 7.25 with KOH). Electrophysiological recordings were amplified using a Multiclamp 700B amplifier (Molecular Devices) filtered at 6 kHz and digitised at 20 kHz with a multifunction I/O device (PCI-6221, National Instruments). Recordings were controlled and analysed offline using Axograph (Axograph, Sydney, Australia). The liquid junction potential (∼9 mV) was not compensated for. The locations of all recorded cells were mapped to coordinates on the Rat Brain Atlas (Paxinos & Watson, 2018).

Recordings were restricted to electrophysiologically-phenotyped glutamatergic projection neurons of the BLA. Projection neurons were distinguished from local circuit GABAergic cells based on their AP waveform and passive electrophysiological properties. Projection neurons have broader APs (>0.7 ms) and smaller fast AHPs (<15 mV) than GABAergic neurons and an input resistance < 150 MΩ. Additionally, projection neurons show frequency-dependent spike broadening, which is absent in GABAergic neurons (Power et al., 2011; Sah et al., 2003). Only cells that had resting potentials more negative than −55 mV, AP amplitudes > 50 mV, and input resistances > 50 MΩ were considered heathy and included in the dataset. Data were also excluded if the series resistance was > 25 MΩ or more than 200 pA was required to maintain the neuron at −65 mV. Series resistance and membrane resistance were calculated from current response to a voltage step from −65 to −70mV using in built routines in Axograph.

### Electrophysiological Protocols and Analysis

For current-clamp protocols, cells were maintained at −65 mV with depolarising or hyperpolarising current injection. The membrane time constant (τ) was determined by fitting an exponential to the voltage response to a small hyperpolarising current.

Intrinsic excitability was calculated as the number of APs evoked in response to a series of 600 ms depolarising current injections (0 to +400 pA, 25 pA increments). The AP threshold, amplitude, half-width, and fast afterhyperpolarisation fAHP were calculated relative to AP threshold and measured from the first AP evoked at rheobase.

The post-burst AHP was evoked by a train of 15 APs evoked by brief (2 ms) somatic injections of depolarising current (2 nA) at a frequency of 50 Hz. The mAHP was the maximal negative amplitude of the post-burst hyperpolarisation within 100 ms of last current injection. The sAHP was defined as the post-burst membrane potential 1000 ms after the AP train.

AHP currents were evoked under a voltage clamp (−50 mV) using a 200 ms depolarising voltage step (+55 mV) and I_AHP**50**_ and sI_AHP**1000**_ currents were measured at 50 ms and 1000 ms after the voltage step, respectively.

### Statistical Analysis

Primary statistical comparisons were performed in Prism 10 (GraphPad, California, USA) using significance levels of α = 0.05. The Shapiro-Wilk test was used to test for normality. For both the behavioural and electrophysical data, between group differences were assessed using a one-way or two-way ANOVA or non-parametric equivalent as appropriate. AP firing data was computed using repeated measures ANOVA. Performance-dependent excitability changes were determined using a simple linear regression comparing average excitability against lever preference. Post-hoc statistical comparisons were performed using the PSY statistical package (UNSW). Data are expressed as mean ± SEM.

## Results

### Behavioural Training

21 rats (9 male, 12 female) underwent behavioural training, with seven allocated to each behavioural group (Punishment, Yoked, and Reward). Sex differences were not examined as this was not the focus of the study. All behavioural groups received lever press training for seven days (Fig. 2A-C). There was an increase in lever presses across days (F_1,18_ = 81.2, p < .001). Rates of responding on R2 were higher than R1 (F_1,18_ = 6.89, p = .017). This was unexpected because the levers had been treated identically. Importantly, however, there were no differences between groups in lever press rates or in the difference in rates of R1 and R2 across days (i.e. no group main effect or group x day interaction) (F_1,18_ = 0.296, p = .59). So, these data show that rats learned to lever press for reward but expressed an initial slight preference for R2 that was equivalent across groups.

**Figure 1:**
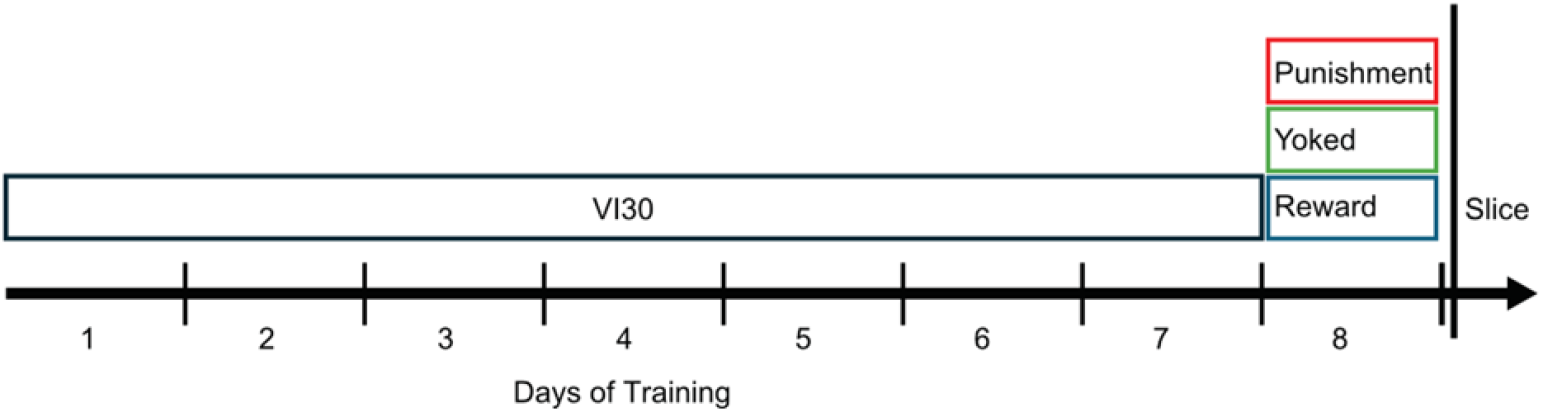
**Timeline of behavioural experiments,** including seven days of variable ratio training, and one day of Punishment, Yoked, or Reward.

**Figure 2:**
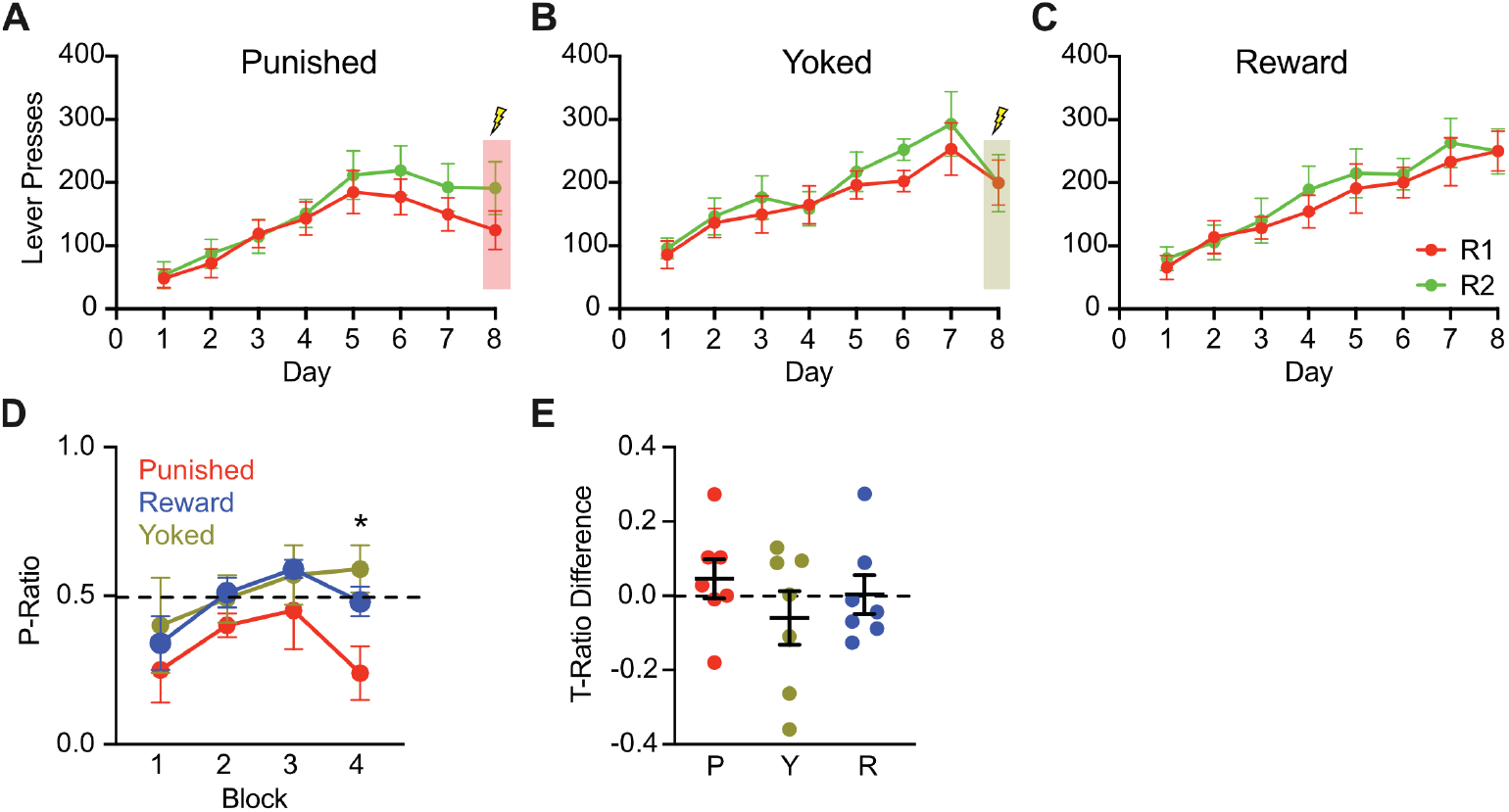
Behavioural Data. (A-C) Lever presses for groups across days 1-8. (D) Day 8 preference ratios (P-Ratio) across four 5-minute time blocks; block 4 P-Ratio was reduced in Punished animals. (E) Overall T-Ratio differences (preference change from day 7 to day 8) for Punished (P), Yoked (Y), and Reward (R). [*] indicates p <0.05.

Footshock was introduced on Day 8. Punished rats received footshocks in response to every tenth press of lever R1 and lever R2 was unpunished. Yoked rats received shocks during presentations of R1 but not R2, independently of their lever pressing. Reward rats received no shocks. The introduction of footshock reduced responding in groups Punished and Yoked (Fig. 2A-C). To evaluate this suppressive effect of footshock and how it affected learning, a P-Ratio was calculated for each 5-minute block of training on Day 8 (Fig. 2D). The P-Ratio represents the rate of responding on the punished lever (R1) relative to the total responding on both levers (R1 and R2). As expected, selective suppression on R1 (P-Ratio < 0.5) emerged with training. Significant group differences in the P-Ratio were observed in the final block of the session (F_2,17_ = 5.61, p = .01). Specifically, Punished rats showed a significant suppression compared to Yoked and Reward rats (F_1,17_ = 10.0, p < .05), with no significant difference in suppression between Yoked and Reward rats (F_1,17_ = 1.18, p > .05). To account for lever preferences, training suppression ratios (T-ratios) were calculated for each lever (T = D8/(D7+D8)) such that lever-pressing on the final session (D8) was measured relative to the previous session (D7). A positive difference between difference between T-Ratios for unpunished and punished levers (T_(unpun)_ − T_(pun)_) > 0 reflects a bias away from the punished lever (Jean-Richard-Dit-Bressel et al., 2019). Although 5 of 7 Punished rats showed positive T-ratio differences, group-level effects were not significant (F2,18 = 0.79, p = .47), likely due to the delayed emergence of suppression within the session (Fig. 2E).

### Ex vivo electrophysiology

To establish whether punishment induces changes to BLA excitability, whole-cell patch-clamp recordings were made from BLA projection neurons in brain slices prepared one hour after the final behavioural session. Seventy-four BLA glutamate neurons were recorded from Punished, Yoked, Reward, and seven Naive rats that had received no behavioural training. Recordings were restricted to neurons within the basal nucleus and from the caudal regions of the BLA, as these areas are implicated in punishment (Jean-Richard-Dit-Bressel & McNally, 2015) (Fig. 3A). There were no group differences in the resting membrane potential (RMP) (F_3,71_ = 0.780, p = .51), membrane time constant (τ) (F_3,71_ = 0.286, p = .84), membrane resistance (Rm) (F_3,71_ = 1.99, p = .12) or series resistance (Rs) (F_3,71_ = 0.644, p = .59) (Table 1).

**Table 1:**
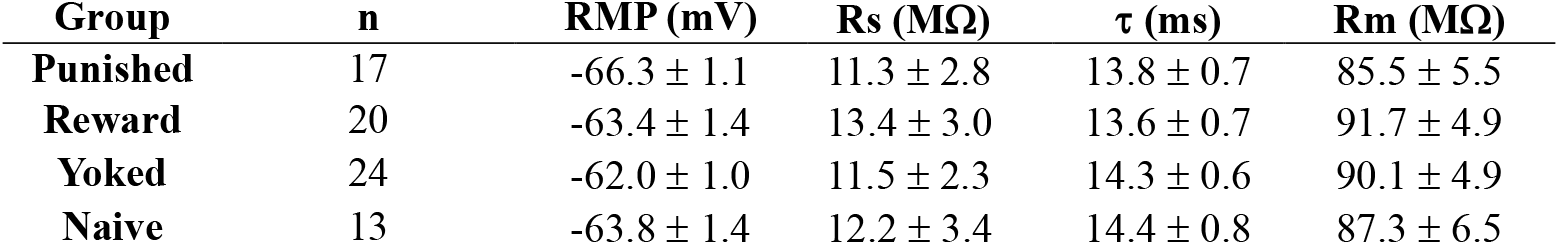
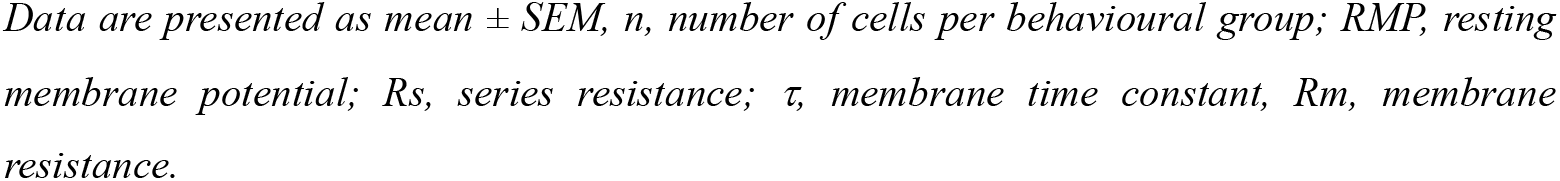
Passive cell properties.

| Group | n | RMP (mV) | Rs (MΩ) | $\tau$ (ms) | Rm (MΩ) |
| --- | --- | --- | --- | --- | --- |
| **Punished** | 17 | -66.3 ± 1.1 | 11.3 ± 2.8 | 13.8 ± 0.7 | 85.5 ± 5.5 |
| **Reward** | 20 | -63.4 ± 1.4 | 13.4 ± 3.0 | 13.6 ± 0.7 | 91.7 ± 4.9 |
| **Yoked** | 24 | -62.0 ± 1.0 | 11.5 ± 2.3 | 14.3 ± 0.6 | 90.1 ± 4.9 |
| **Naive** | 13 | -63.8 ± 1.4 | 12.2 ± 3.4 | 14.4 ± 0.8 | 87.3 ± 6.5 |

*Data are presented as mean ± SEM, n, number of cells per behavioural group; RMP, resting membrane potential; $R_s$ , series resistance; $\tau$ , membrane time constant, $R_m$ , membrane resistance.*

**Figure 3:**
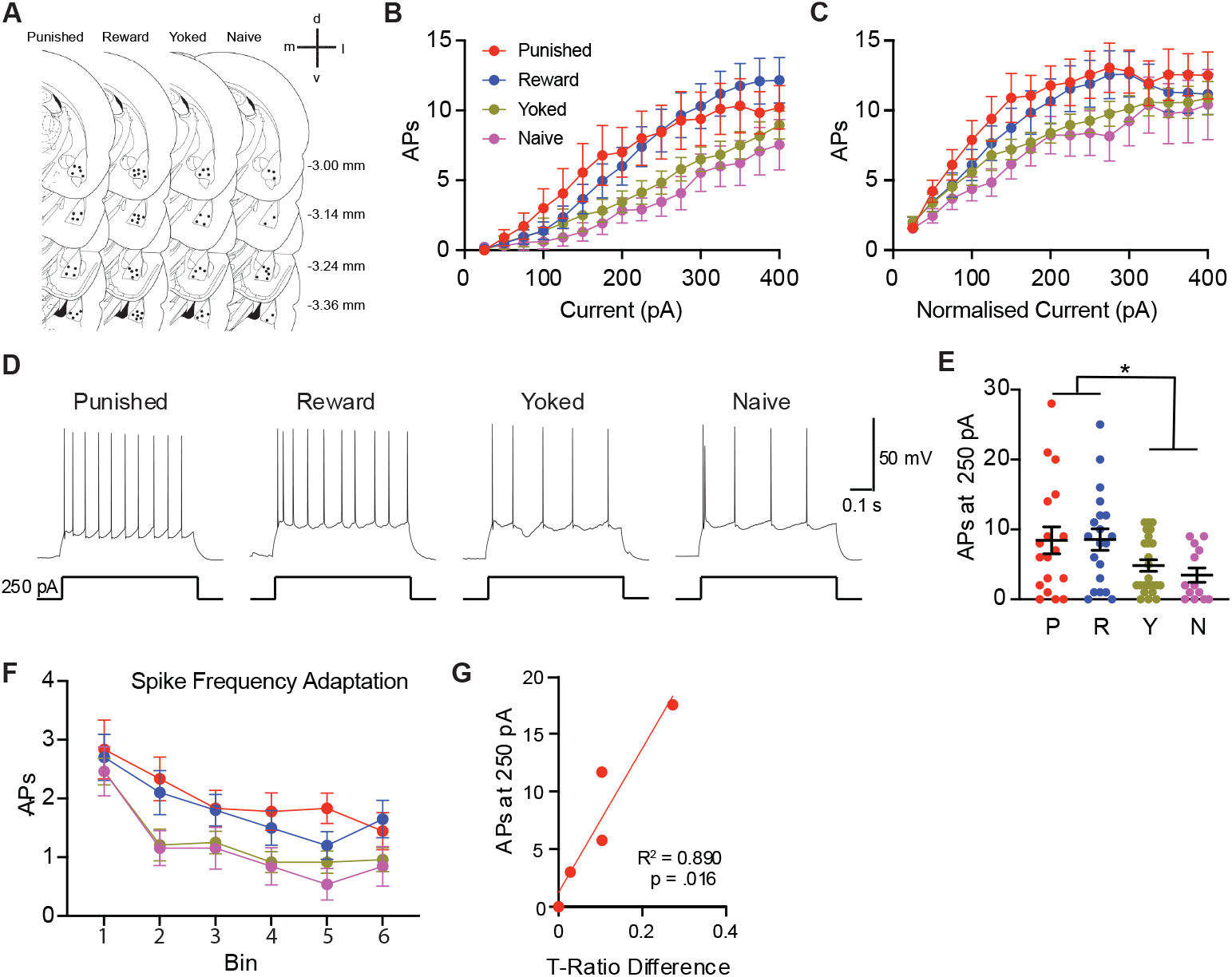
Learning-Associated Intrinsic Excitability Changes. Anatomical distribution of recorded BLA neurons (A). Estimated distance from bregma shown to the right. APs as a function of raw current injection for neurons from Punished (P), Reward (R), Yoked (Y), and Naive (N) rats. (B). APs normalised to rheobase current, with increasing firing at higher current amplitudes (C). Spikes evoked by 250 pA, neurons from showing that Yoked and Naive animals had reduced firing compared to Punished and Reward (D-E). Number of APs evoked by a 600 ms current injection (100 pA above rheobase) plotted at 100 ms intervals (F). Regressions performed between training ratio (T-Ratio) differences and mean spikes evoked by 250 pA mean per rat from firing properties data for Punished rats (G), showing that BLA excitability predicted behavioural bias towards the unpunished lever. [*] indicates p < 0.05

### Effect of Training on Intrinsic Excitability

We next examined whether training altered intrinsic neuronal excitability by examining the number of APs evoked by a series of depolarising 600 ms somatic current injections (Fig. 3B). Two-way repeated measures ANOVA revealed an increase in AP firing as a function of injected current (F_14_,_994_ = 60.0, p < .0001) and significant interaction between current and behavioural group (F_42, 994_ = 1.7, p = .0028), but no overall effect of behavioural group (F_3, 71_ = 2.7, p = .05). Similar effects were observed when measured relative to the minimal current amplitude to evoke APs (rheobase) with an increase in AP firing as a function of injected current (F_19, 1331_ = 40.1, p < .0001), no behavioural groups difference (F_3, 71_ = 1.37, p = .26), and a significant interaction between current and behaviour group (F_21,493_ =1.8, p = .013; Fig. 3C). To further examine these changes, we compared the number of APs evoked by a 250-pA current step (Fig. D-E). This stimulation intensity was chosen to avoid a floor effect on spike counts (likely at lower current injections) and to minimise depolarisation block (likely at higher current injections) (Bianchi et al., 2012). The number of spikes evoked by a 250-pA current step differed between groups (F_3,71_ = 3.13, p = .031) confirming the initial finding. Follow up analyses revealed greater spike counts at 250 pA for the Punished and Reward groups versus Yoked and Naive groups (F_1,71_=9.37, p < .05). There was no difference between Punished and Reward groups (F_1,71_ = 0.003, p > .05) or between Yoked and Naive groups (F_1,71_ = 0.44, p > .05). Taken together these findings show two things. First, instrumental training in the form of either Punishment or Reward is associated with an elevation in the excitability of BLA projection neurons. Second, non-contingent shock delivery arrests this excitability increase and returns the excitability of BLA projection neurons to a Naive state.

Learning-induced changes to the input-output relationship often results from a reduction in the spike frequency adaptation normally characteristic of BLA projection neurons and hippocampal pyramidal neurons (Motanis et al., 2014; Oh & Disterhoft, 2015; Sehgal et al., 2023; Sehgal et al., 2014). To assess spike frequency adaptation, we examined the number of APs generated at 100 ms intervals along the current step (100 pA above rheobase) (Fig. 3F). As expected, the number of APs decreased across the time intervals (F_2.329,165.3_ = 38.0, p < .0001) but no behavioural group difference (F_3,71_ = 2.45, p = .07) or interaction between behavioural group and time interval (F_15,355_ = 1.11, p = .35) was detected.

### Intrinsic Excitability and Learning Performance

The findings thus far show that instrumental training is associated with an elevation in the excitability of BLA projection neurons. However, these data do not show any difference between Punishment and Reward. Therefore, it is unclear whether and how these intrinsic excitability changes relate to punishment learning specifically versus the instrumental contingencies shared across groups, especially the Punished and Reward groups (lever presses delivered pellets for both groups). To address this, we next examined whether changes in BLA firing rates predicted differences in responding on the two levers. Linear regressions were performed comparing the differences in lever press training ratios against the mean spike count evoked by 250 pA for the recorded cells from each rat. There was no significant relationship between behavioural bias (i.e. differences in lever press training ratios) and the mean spikes evoked by 250 pA for the Punished (R^2^ = 0.246, p = .26), Yoked (R^2^ = 0.00270, p = .91), or Reward groups (R^2^ = 0.0001, p = .98). However, as mentioned above, there were individual differences in the behavioural bias (T-Ratio) within the Punished group. When the correlation analysis was limited to animals that exhibited behavioural bias from punishment learning (T-Ratio > 0), increased excitability was found to correlate with the behavioural bias (R^2^ = 0.890, p = 0.016; n = 5; Fig. 3G).

### Effect of training on the AP and AHP

To determine the mechanisms underpinning differences in firing, AP waveform parameters were measured under current clamp (Fig. 4). One-way ANOVAs showed no between group differences for fAHP (F_3,71_ = 1.19, p = .41), AP amplitude (F_3,71_ = 1.40, p = .25) or AP threshold (F_3,71_ = 2.64, p = .06) but did show a significant difference in AP half-width (F_3,71_ = 3.91, p = .012) (Fig. 4B-F). Post-hoc analysis showed that groups that had received footshock (i.e. Punished, Yoked) had significantly lower AP half-widths compared to the non-footshock groups (i.e. Reward, Naive) (F_1,71_ = 11.3, p < .05). There was no difference between groups that had received footshock (i.e. Punished, Yoked) (F_1,71_ = 0.07, p > .05) or between non-footshock groups (i.e. Reward, Naive) (F_1,71_ = 1.44, p > .05). To test for the impact of learning on AP half-width, punished animals were divided into two groups: *learners* (those with T-ratios > 0, indicating a preference for the punished lever) and *non-learners* (T-ratios < 0, indicating a preference for the unpunished lever). No significant differences in AP half-width were observed between the two groups (unpaired t-test: t(15) = 1.44, p = 0.13).

**Figure 4:**
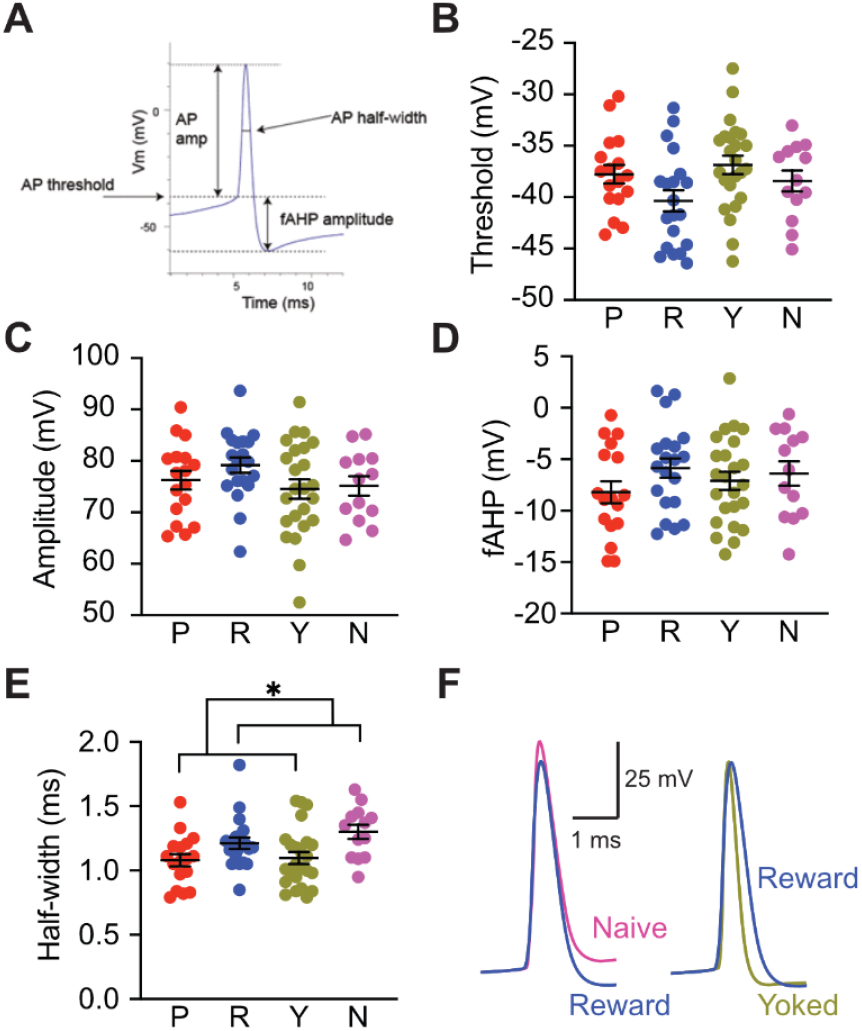
Effect of conditioning on AP Waveform. Diagrammatic representation of AP waveform measures. (A). Summary data showing AP threshold (B), AP amplitude (C), fAHP amplitude (D), and AP half-width (E) for neurons from Punished (P), Reward (R), Yoked (Y), and Naive (N) rats. APs from Punished and Yoked subjects were narrower than Reward and Naive. Representative APs from Naive, Reward, and Yoked neurons (F). [*] indicates p < 0.05.

In addition to examining AP waveform, we also tested the amplitude of the medium AHP (mAHP) and slow AHP (sAHP) that follow a burst of APs (Fig. 5). No between group differences were observed for the mAHP (F_3,71_ = 1.39, p = .14) or sAHP measured 1 s after the AP train (F_3,71_ = 1.71, p = .17) (Fig. 5B-C). We also examined the currents underlying the post-burst AHP which are primarily mediated by two temporally overlapping calcium-activated potassium currents; the SK-mediated (I_AHP_) and the slower apamin-insensitive sI_AHP_ (Power et al., 2011; Power & Sah, 2008). Under voltage clamp (Vm = −50 mV), group differences were observed in the AHP current 50 ms after the voltage step I_AHP50_ mediated by both currents (F_3,71_ = 7.86, p = .0001), but not the current 1000 ms after the voltage step I_AHP1000_ (F_3,71_ = 1.16, p = .33), to which only the sI_AHP_ contributes (Fig. 5E-F). Post-hoc analysis comparing groups that had received footshock (i.e. Punished, Yoked) had significantly larger I_AHP50_ compared to the non-footshock groups (i.e. Reward, Naive) (F_1,71_ = 15.4, p < .05) but no difference between the groups that had received footshock (i.e. Punished, Yoked) (F_1,71_ = 0.63, p > .05) or between the non-footshock groups (i.e. Reward, Naive) (F_1,71_ = 3.63, p > .05).

**Figure 5:**
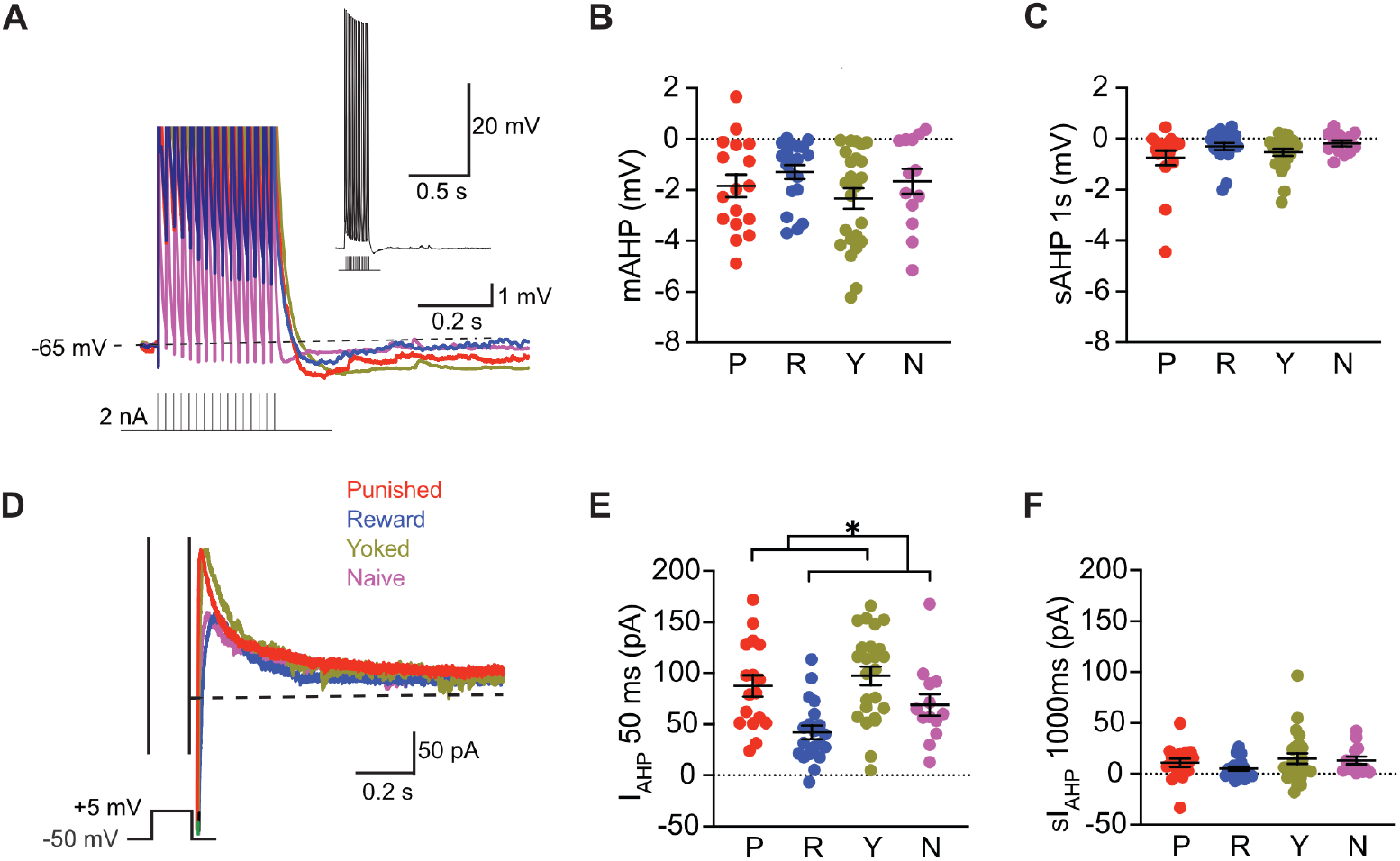
AHP Differences Across Behavioural Groups. (A) Representative voltage response a 300 ms 50 Hz train of 2 nA 2 ms current pulses shown above (inset) the representative responses from Punished (P), Reward (R), Yokes (Y), and Naive (N) neurons shown on an expanded voltage scale. Summary data showing the peak mAHP (B) and slowAHP 1s after the train (C). Representative currents evoked by a 200 ms 55 mV voltage step (D). Summary data showing the AHP current measured 50 ms, I_AHP_50 ms (E) and 1000 ms, sI_AHP_1000 ms (F) after the voltage step. The I_AHP**50**_ amplitudes was larger in Punished and Yoked animals compared to Reward and Naive (D-E). [*] indicates p < .05

## Discussion

This study examined whether instrumental learning induces changes in BLA projection neuron intrinsic excitability. We found that both reward and punishment training increased intrinsic excitability relative to yoked and naive controls. Notably, excitability was positively correlated with behavioural performance on the punishment task, suggesting a functional relationship between neuronal excitability and learning. In addition, we observed group differences in AP half-width and the current underlying the post-burst AHP, pointing to distinct biophysical adaptations following different forms of learning.

Previous studies have shown that Pavlovian fear conditioning enhances BLA projection neuron excitability (Sehgal et al., 2023; Sehgal et al., 2014). Our findings extend this work by showing that instrumental learning, both appetitive and aversive, similarly increases intrinsic excitability. Such increases are thought to facilitate integration of neurons into functional learning circuits and support synaptic integration and long-term potentiation (Choi et al., 2021; Josselyn & Frankland, 2018; Park et al., 2023; Power et al., 2011; Sah & Bekkers, 1996). These results suggest that heightened intrinsic excitability may be a general feature of BLA neurons recruited into action-outcome learning, regardless of outcome valence.

Interpretation of punishment effects is confounded by punishment being layered on top of instrumental reward learning. While punishment and reward groups did not differ significantly in AP output to current injections, the correlation between excitability and punishment performance, along with lower excitability in the Yoked group - which received identical shock exposure in a non-contingent manner - suggests that the instrumental punishment contingency plays a critical role in maintaining or enhancing excitability. This finding raises the possibility that punishment-specific learning processes can preserve or augment the excitability increases initiated during reward training.

Surprisingly, intrinsic excitability was not enhanced in the Yoked group which received non-contingent footshocks akin to Pavlovian fear conditioning. In fact, excitability in the Yoked group resembled that of naive controls and was lower than in both instrumental groups. This result contrasts with prior studies showing fear-induced increases in BLA excitability (Sehgal et al., 2023; Sehgal et al., 2014), though methodological differences may account for the discrepancy. While those studies targeted neurons in the lateral amygdala, our recordings sampled from the basal nucleus, which has distinct projection targets and functional roles (Beyeler et al., 2018; Namburi et al., 2015). Furthermore, our task was conducted in a reward-associated context in the absence of an explicit conditioned stimulus. Supporting this finding, (Butler et al., 2018) reported no change in intrinsic excitability following contextual fear conditioning when sampling broadly across the BLA. Together, these results suggest that contextual fear conditioning, when superimposed on instrumental training or delivered in a minimally salient context, may not engage the same excitability-enhancing mechanisms observed in classical cued fear paradigms.

Intrinsic excitability is regulated by numerous ion channels, and learning-related increases are often associated with reduced activity of the slow calcium-activated potassium current (sI_AHP_) underlying the post-burst AHP (McKay et al., 2013; Sehgal et al., 2014; Titley et al., 2020). However, we did not observe significant learning-related differences in the post-burst AHP or the isolated sI_AHP_. This may be due to the small amplitude and thus limited resolution of our recorded AHPs, likely resulting from use of gluconate rather than MeSO4 as the primary internal solution anion (Kaczorowski et al., 2007; Zhang et al., 1994).

We did detect group differences in the AHP current 50 ms after the voltage step (I_AHP_50), which reflects contributions from both apamin-sensitive SK channels and apamin-insensitive sI_AHP_ components (Faber & Sah, 2002; Power & Sah, 2008). Post hoc analysis showed that I_AHP_50 was larger in animals exposed to shock—both Pavlovian and instrumental - compared to non-shocked groups, suggesting that shock exposure independently modulates AHP-related currents, although its relationship to overall excitability remains unclear.

Group differences in AP half-width also emerged, with narrower spikes observed in shock-exposed animals (Punished and Yoked) compared to non-shocked animals (Reward and Naive). Although AP half-width has been reported to increase following extinction of Pavlovian fear (Senn et al., 2014), it is often inversely correlated with firing frequency (Faber & Sah, 2002). In our case, the narrowing of spikes may reflect a distinct biophysical adaptation linked to shock exposure rather than learning per se.

Together, these findings show that successful instrumental punishment learning increases BLA projection neuron excitability, resembling the effects of reward rather than fear learning. In contrast, Pavlovian shock presentation appears to halt or reverse these excitability increases, returning neurons to a state resembling baseline. Exposure to shock, whether contingent or non-contingent, also induced distinct biophysical changes - including increased I_AHP_ amplitude and narrower AP half-widths - suggesting that aversive stimuli modulate membrane properties independently of learning contingencies. These results clarify that punishment and fear are encoded differently at the cellular level within the BLA, with punishment-associated excitability more closely resembling reward-driven plasticity.

Finally, our findings may be relevant to understanding psychiatric disorders marked by altered punishment sensitivity, such as major depression and conduct disorder. Punishment-specific excitability changes in the BLA may reflect a distinct neural coding scheme, separate from that recruited by fear, and may help explain why maladaptive behaviours persist even in the face of negative consequences.

### Limitations and Future Directions

It is important to note that several of the analyses reported here were conducted *post hoc*. While these analyses revealed meaningful group differences and informed interpretations of the data, they were not part of our *a priori* experimental plan. As such, the findings - particularly those related to AP half-width and I_AHP_50 - should be considered exploratory. Future studies will be critical to confirm these observations and determine their mechanistic basis. Additionally, employing methods with greater sensitivity to AHP components - such as alternative internal solutions or sharp electrode recordings - may help clarify the role of specific ionic currents in learning-related excitability changes.

## Acknowledgments

Author Contributions: Conceptualisation: JMP, GPM, ETW; Methodology: JMP, GPM, ETW, PJRdB, JOYY; Investigation: ETW, JOYY; Writing - Original Draft: ETW, JMP, GPM. Writing - Review & Editing: All authors. Funding Acquisition: JMP, GPM. This work was supported by a Research Training Program Scholarship, and the National Health and Medical Research Council (Ideas GNT2011633)

